# Accurate inference of genome-wide spatial expression with iSpatial

**DOI:** 10.1101/2022.05.23.493144

**Authors:** Chao Zhang, Renchao Chen, Yi Zhang

## Abstract

Spatially resolved transcriptomic analyses can reveal molecular insights underlying tissue structure and context-dependent cell-cell or cell-environment interaction. Due to the current technical limitation, obtaining genome-wide spatial transcriptome at single-cell resolution is challenging. Here we developed a new algorithm named iSpatial to derive spatial pattern of the entire transcriptome by integrating spatial transcriptomic and single-cell RNA-seq datasets. Compared to other existing methods, iSpatial has higher accuracy in predicting gene expression and their spatial distribution. Furthermore, it reduces false-positive and false-negative signals in the original datasets. By testing iSpatial with multiple spatial transcriptomic datasets, we demonstrate its wide applicability to datasets from different tissues and by different techniques. Thus, we innovated a computational approach to reveal spatial organization of the entire transcriptome at single cell resolution without the need of new technology development. With numerous high quality datasets available in the public domain, iSpatial provides a unique way for understanding the structure, function of complex tissues and disease processes.

**HIGHLIGHTS:**

- iSpatial infers genome-wide spatial gene expression pattern by integrating spatial transcriptomic and scRNA-seq data
- iSpatial outperforms existing approaches in inferring spatial gene expression patterns
- iSpatial reduces false-positive/negative signals of the original spatial transcriptome
- iSpatial is applicable to spatial transcriptomic datasets from different tissues and techniques

## INTRODUCTION

In the past decade, single-cell RNA-sequencing (scRNA-seq) has transformed our understanding on the cellular heterogeneity of various tissues/organs in multicellular organisms (*1–4*). With current scRNA-seq techniques, obtaining whole transcriptomic profiles of tens to hundreds of thousands of single cells has become a routine. However, most high-throughput scRNA-seq methods use dissociated cells and consequently the spatial information of the analyzed cells is lost, which prevents directly connecting the molecular feature of the analyzed cell types to their anatomic and functional features. On the other hand, the development of spatially resolved transcriptomic assays have enabled the transcripts/cells location analysis in the tissue context, which has the potential to reveal how single-cell gene activity orchestrates the structure and function of complex tissues in health and disease (*5*).

In the past few years, different methods for spatial transcriptomic (ST) assays have been developed (*6–13*). Ideally, the spatial transcriptome data should provide genome-wide and spatially resolved expression measurements at single-cell resolution. However, due to technical limitations, either spatial resolution or gene coverage is compromised in most ST assays. For example, *in situ* capture and sequencing-based techniques are able to capture any mRNA molecules without pre-knowledge, but the spatial resolution is not at single-cell level (*6, 14*). On the other hand, *in situ* sequencing and FISH-based mRNA measurement can achieve cellular or subcellular resolution, but most such assays are limited with their throughput on genes that can be detected (usually 30 −500), and require pre-knowledge for probe design (*7–9*).

With the rapid development of scRNA-seq and ST technologies, new bioinformatic tools have been developed to overcome the challenges in single-cell and spatial genomic data analysis (*15, 16*). Notably, by integrating scRNA-seq and spatial resolved profiling data, recent computational methods have leveraged the strength of different datasets and revealed information that otherwise cannot be obtained from a single experimental paradigm. For example, when the corresponding scRNA-seq is available, SpatialDWLS and RCTD could perform deconvolution on the “low-resolution” spatial dataset to estimate the cell type/proportion in each spatially resolved spot (*17, 18*). On the other hand, Tangram and Cell2location could predict the spatial location of molecularly defined cell types (from scRNA-seq) based on ST data (*19, 20*). Seurat and Liger could impute transcriptome-wide spatial expression by integrating with corresponding scRNA-seq data and transferring the expression data to spatial transcriptome (*21, 22*). Despite these existing tools, inferring expression patterns of all genes at high spatial resolution by integrating scRNA-seq and ST data is not straightforward, and different approaches have variable performance when applied to different datasets. Since the performance of this task is critical for downstream spatial analysis, a robust and convenient tool for spatial pattern prediction is highly desirable.

Here, we present iSpatial, an R-based bioinformatic tool that integrates scRNA-seq and ST profiling data to infer the expression pattern of each gene at high spatial resolution. We show that iSpatial outperforms existing approaches on its accuracy and it can also reduce false-positive and false-negative signals in the original data. By applying iSpatial to datasets from different tissues (hippocampus, cortex, striatum and liver) and generated with different techniques (Slide-seq, MERFISH, STARmap), we have revealed both known and novel spatial expression patterns in each dataset, indicating iSpatial is broadly applicable for analyzing different ST datasets. Collectively, our analyses demonstrate that iSpatial is a useful tool for resolving transcriptom-wide spatial expression patterns at single cell resolution in complex tissues.

## RESULTS

### Overview of iSpatial

The FISH and *in situ* sequencing-based ST techniques, such as MERFISH (*7*), seqFISH (*8*) and STARmap (*9*), can simultaneously reveal gene expression and location at single-cell resolution, but with limited pre-defined gene targets (**Fig. 1A**, left). On the other hand, scRNA-seq can unbiasedly profile the whole transcriptome, but without providing spatial information (**Fig. 1A**, middle). We reasoned that by integrating the single-cell gene expression profiles (the gene by cell matrices) of the two methods, the missing information of non-targeted genes in each spatially profiled cells could be inferred based on scRNA-seq data, resulting in genome-wide spatial expression information of the profiled cells (**Fig. 1A**, right).

**Fig. 1:**
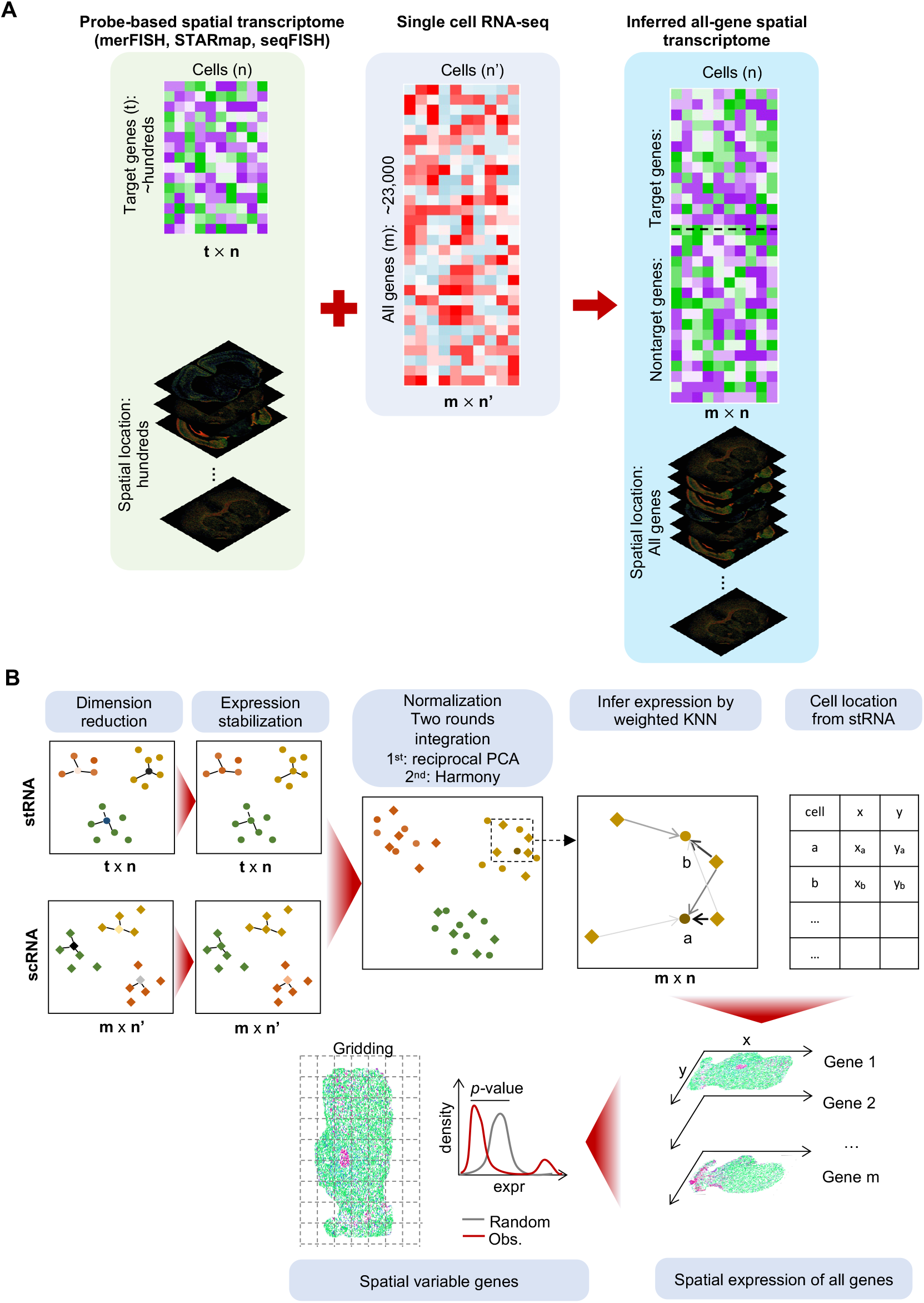
Overview of iSpatial. **A**, A diagram showing the rationale of iSpatial. The probe-based spatial transcriptome includes *t* × *n* (*t,* genes; *n,* cells) expression matrix and location of each cell. After integrating with *m* × *n*′ scRNA-seq data, iSpatial infers the genome-wide transcriptional expression of all *n* cells. **B**, The iSpatial pipeline is consisted of: 1) Dimension reduction and expression stabilization; 2) Expression normalization; 3) Inferring transcriptional expression; and 4) Downstream analysis such as spatial variable gene detection.

To this end, we first performed dimension reduction on scRNA-seq and spatial profiling data separately, followed by expression stabilization, which removes potential false-positive and false-negative expression values based on the expression level of adjacent cells in PCA space. The two datasets were then normalized and embedded into a common space with two sequential rounds of integration: first by reciprocal PCA (*21*) and then through Harmony (*23*). Based on the common embedding, the expression value of each gene in each cell of the ST dataset is then inferred using a weight k-nearest neighbors (KNN) model. Since the physical locations of these cells have been resolved in spatial profiling, the results represent a new single-cell gene expression profile with both genome-wide coverage and single-cell spatial resolution, which could be used for downstream analyses including identification of spatially variable genes (SVGs) (**Fig. 1B**).

### iSpatial outperforms existing tools in its accuracy on predicting spatial expression pattern

To evaluate the performance of iSpatial and compare it with existing tools, we used a mouse hippocampal dataset generated from Slide-seq V2 (*24*). Since this dataset includes the spatial expression of all genes, it can be used for evaluating the prediction performance (**Fig. 2A**). Specifically, we divided the dataset into the training and validation groups, which contain 3,000 and ∼20,000 genes, respectively. The training data group (mimic a ST dataset) was integrated with a scRNA-seq dataset covering the same brain region (hippocampus) (*1, 25*) (but by a different method) to infer the expression level and spatial patterns of the genes in the validation data group. By comparing the inferred expression patterns with the “truth” determined by Slide-seq (validation data group), we found that iSpatial could predict the spatial expression pattern with high accuracy. For example, iSpatial inferred the expression of *Atp2b1*, *Prox1* and *Fibcd1*, which were not included in the training data group, across the entire hippocampus, dentate gyrus (DG), and CA1, respectively, consistent with the Slide-seq validation data and ISH results from Allen Brain Atlas (ABA) (*25*) (**Fig. 2B**). Interestingly, we found that iSpatial could “enhance” the signals not well detected in the original data. For example, *Slit1*, *Tspan18*, *Efnb2*, *Car12* and others were barely detectable in hippocampal cells by Slide-seq, thus difficult to determine their spatial pattern. With iSpatial, the expression of these genes was clearly visible, thus their spatial pattern could be clearly recognized. This is unlikely an artifact of imputation, as the spatial expression inferred by iSpatial is consistent with that of the ABA data (**Fig. 2C and fig. S1A**).

**Fig. 2:**
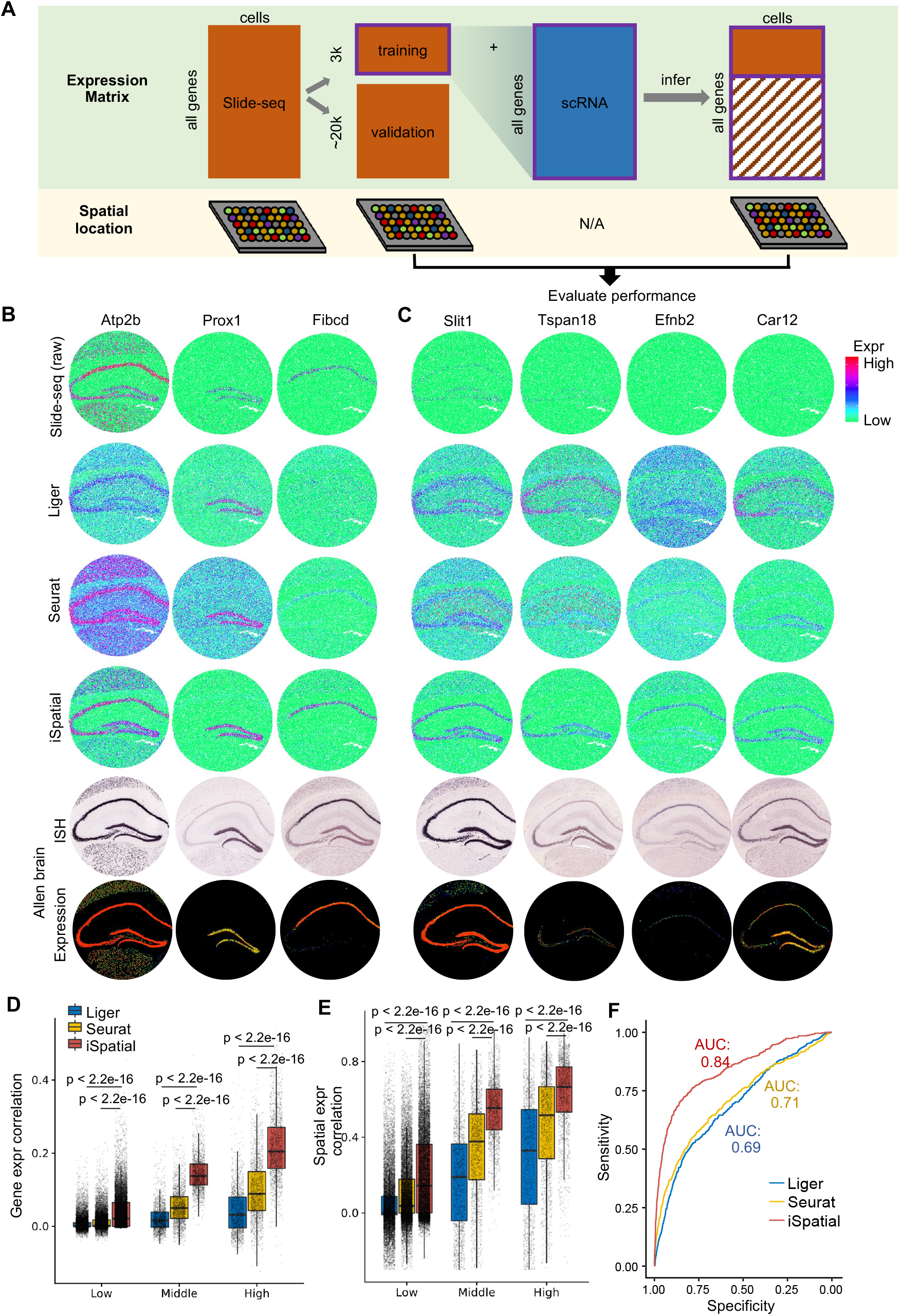
Benchmarking the performance of iSpatial in inferring genome-wide spatial transriptome. **A**, A graphical illustration of the evaluation procedure. **B**, Representative examples showing the performance of Liger, Seurat and iSpatial in inferring spatial transcriptome. **C**, Representative examples of inferred expression of genes barely detectable in raw Slide-seq data. **D**, The gene expression correlation between Slide-seq raw data and Liger, Seurat, or iSpatial inferred data. The validation genes are divided into 3 groups based on their expression levels. Two-side Wilcoxon Rank Sum test was used. **E**, The spatial expression correlation between inferred and raw Slide-seq data. Two-side Wilcoxon Rank Sum test was used. **F**, The ROC curves comparing the prediction power of spatial variable genes by Liger, Seurat, and iSpatial. AUCs (Area under the curves) are indicated.

We further compared the performance of iSpatial with another two existing tools, Liger (*22*) and Seurat (*21*), on the same task using the Slide-seq dataset. Although these two methods could also infer the expression patterns of genes not included in the training data group, compared with iSpatial, the spatial patterns obtained from Liger and Seurat were more ambiguous with higher background in general (**Fig. 2B, 2C and fig. S1A**). To quantitatively benchmark these different methods, we calculated the expression and spatial correlation between Slide-seq data (regarded as ground truth) and inferred results from iSpatial, Liger or Seurat on each gene of the validation dataset. The results showed that the accuracy of prediction is positively correlated with the gene expression level in general, but iSpatial exhibited significantly higher accuracy than the other methods across all gene groups with different expression levels (**Fig. 2D and 2E**). Furthermore, when compared the SVGs identified from the original Slide-seq data with those identified from inferred data of different methods (**fig. S1B**), we found that iSpatial has the highest prediction power with area under the curve (AUC) over 0.84 on SVGs among the three methods (**Fig. 2F**). Collectively, these analyses indicate that iSpatial outperforms existing tools in terms of accuracy on predicting spatial expression pattern.

### iSpatial is broadly applicable to different spatial transcriptomic datasets

After validating the performance of iSpatial with Slide-seq data, we further tested whether it could be applied to other ST datasets from different tissues generated with different techniques. To this end, we first used iSpatial to analyze a STARmap dataset which covered the primary visual cortex (V1) of mouse brain (*9*) (**Fig. 3A**). Although the original STARmap data only included 1,020 gene targets, iSpatial successfully inferred the expression of over 20,000 genes by integrating a single cell smart-seq dataset from ABA (*26*) (**Fig. 3B**). Importantly, the spatial expression patterns of genes not included in the original STARmap data could be faithfully inferred by iSpatial. For example, the layer-specific expression of a number of genes were accurately predicted as evidenced by its similarity to that of the ABA ISH results (**Fig. 3C**). Notably, iSpatial not only correctly predicted the layer distribution of non-targeted genes, but also detected the expression variation of certain genes based on their spatial locations. For example, it predicted: 1) a high-to-low gradient of *Pvrl3* across upper cortical layers, and 2) strong expression of *Serinc2* and *Col5a1* in cortical layer VI but relatively weak expression in upper layer V, both of which were confirmed by ISH data (**Fig. 3C**).

**Fig. 3:**
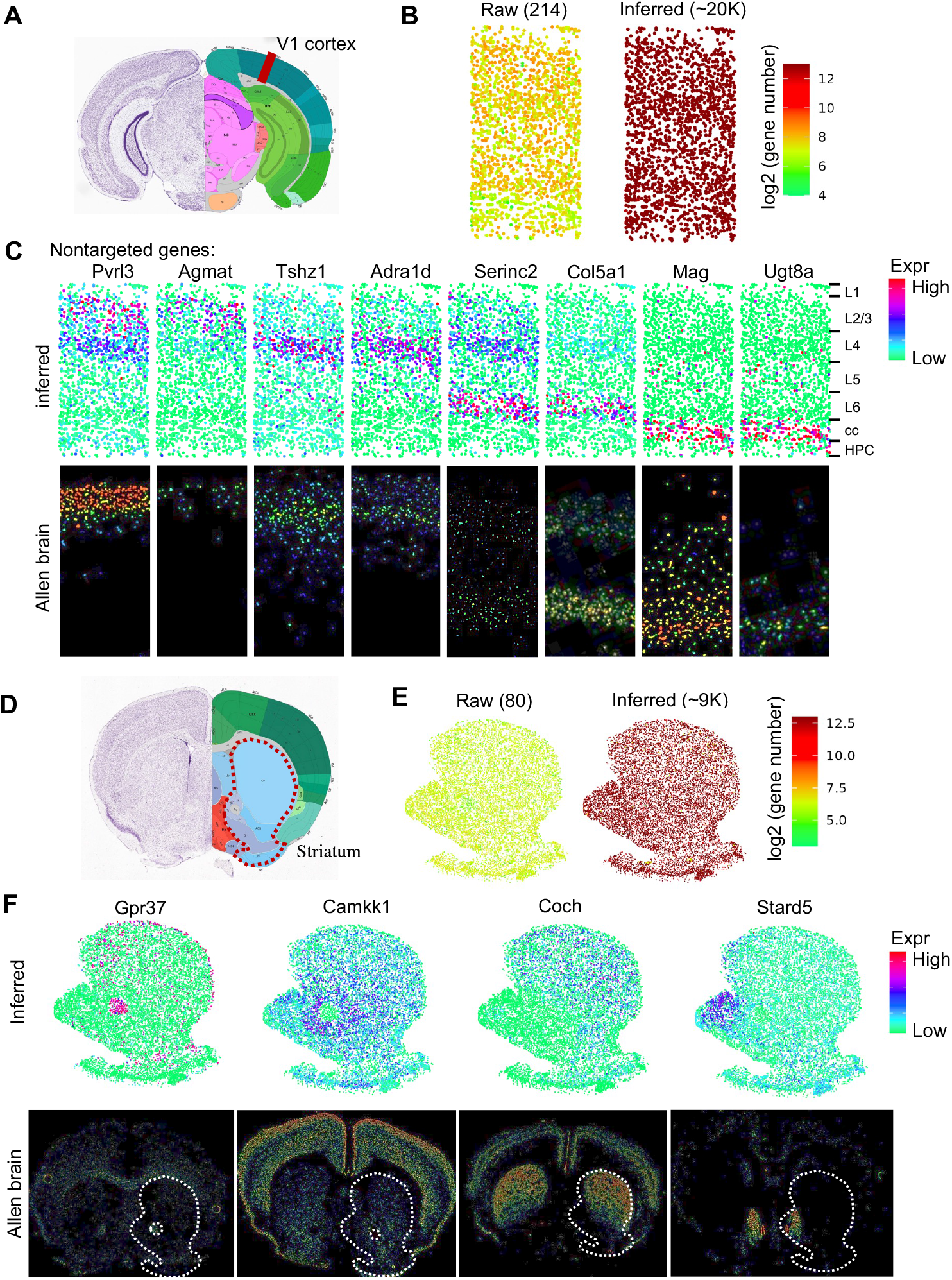
iSpatial accurately infers the genome-wide spatial transcriptomes in mouse cortex and striatum. **A**, Schematic of the anatomic region of mouse V1 cortex. **B**, The numbers of detectable genes in each cell in raw STARmap (left) and after inferring by iSpatial (right). The mean number of detected genes in each cell are shown in brackets. **C**, Inferred layer expression patterns of representative genes not targeted in the original STARmap library compared with the ISH data from the ABA. **D**, Schematic of the anatomic region of mouse striatum. **E**, The numbers of detectable genes in each cell in raw MERFISH (left) and after inferring by iSpatial (right). **F**, Inferred spatial expression patterns of representative genes not targeted in the original MERFISH library compared with the ISH data from the ABA.

In additional to the STARmap dataset, we analyzed a recently published MERFISH dataset of mouse striatum (*27*) with iSpatial (**Fig. 3D**). The original MERFISH dataset contained 253 target genes which allowed the identification of 9 major cell types in striatum (**fig. S2A**), with 175 target genes exhibit significant enrichment in certain cell types (markers) (**fig. S2B**). By integrating this dataset with corresponding scRNA-seq data, iSpatial could infer the expression and location of ∼9,000 genes at single-cell resolution (**Fig. 3E and fig. S2C**), with over 2,200 genes identified as markers for different cell types (**fig. S2D and S2E**). Importantly, the spatial patterns of inferred genes were largely consistent with those determined by ISH. For example, *Gpr37* was highly enriched in the anterior commissure (AC), *Coch* formed a high-to-low gradient from the dorsolateral to the ventromedial striatum, and *Stard5* was specifically expressed in the medial nucleus accumbens (NAc), all these iSpatial inferred expression patterns are consistent with the results from ABA (**Fig. 3F**).

To globally evaluate the predication accuracy, we adopted a 10-fold cross validation approach and found that iSpatial showed significantly higher accuracy than Liger and Seurat in predicting the expression and spatial location of genes in both datasets (**fig. S3**). Collectively, these results demonstrated the capacity of iSpatial in predicting the expression and location of genes using ST dataset generated from different tissues with different techniques.

### iSpatial reduces false-positive and false-negative signals from spatial transcriptome

Although imaging-based transcriptomic assays have a higher detection efficiency when compared to that of sequencing-based methods, their performance is highly variable depending on the specific gene probes. For example, some transcripts are too short to be targeted by enough probes, which may lead to false-negative. On the other hand, some other genes may have close homologs which are difficult to distinguish with hybridization, leading to false-positive. We hypothesized that iSpatial could reduce these false signals by giving higher weights to cells of scRNA-seq when performing expression prediction, which were insensitive to gene length and could also unambiguously distinguish similar transcripts based on sequence differences. To test this hypothesis, we first compared the expression pattern of some well-establish cell type markers on the UMAP between the original STARmap data and iSpatial inferred data. We found that although these cell type-specific markers exhibited high enrichment in corresponding cell types, there were often false-positive signal in other cell types when analyzed by STARmap (**Fig. 4A**, upper panels, *Slc17a7*, *Gad1*, *Plp1* and *Cldn5*). In some cases, the expected expression patterns were not observed, likely due to false-negative (**Fig. 4A**, upper panels, *Aqp4*). Consistent with our hypothesis, iSpatial could remove most false-positive signals from irrelevant cell types, without affecting the true positive signals (**Fig. 4A**, lower panels, *Slc17a7*, *Gad1*, *Plp1* and *Cldn5*). Furthermore, iSpatial successfully inferred gene expression pattern that was missed in the original STARmap analysis (**Fig. 4A**, lower panels, *Aqp4*), suggesting that iSpatial can also reduce false-negative results. Based on these findings, we further asked whether iSpatial could infer spatial patterns which were not well detected by STARmap. Indeed, we found a number of genes, whose expected layer-specific expression patterns were not detected in the original STARmap analysis. For example, *Nov*, *Rorb*, *Rspo1*, *Fezf2*, *Foxp2* are established markers of different cortical layers. However, STARmap only detected sparse signals or even failed to detect real signals of these genes across all cortical layers (**Fig. 4B**). In contrast, iSpatial accurately captured the layer-specific expression patterns of these genes that were also detected by ISH (**Fig. 4B**).

**Fig. 4:**
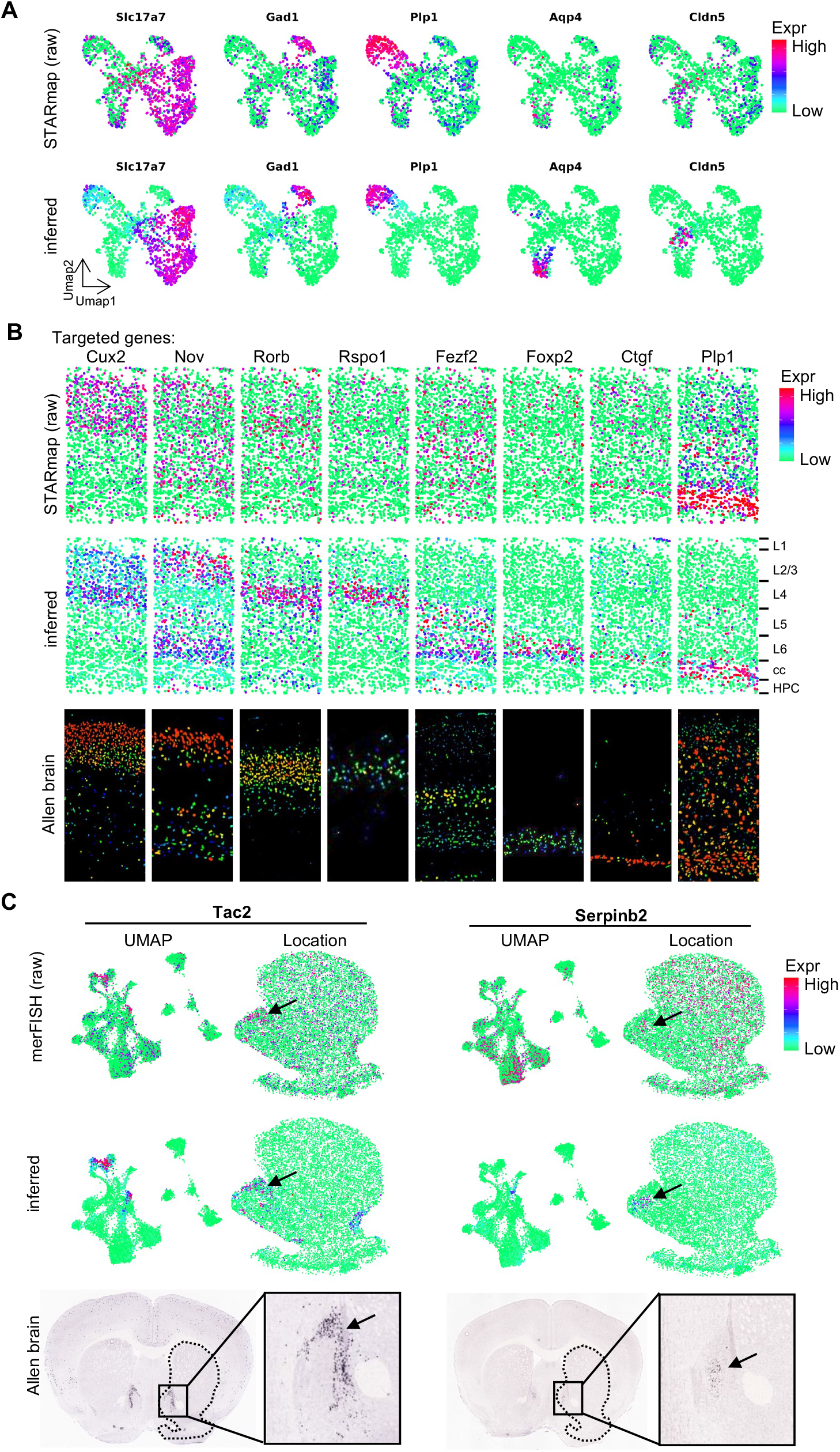
iSpatial can reduce false-positive and false-negative signals in the original spatial transcriptomic data. **A**, UMAPs showing the expression levels of representative cell type markers in raw STARmap (top panels) and iSpatial inferred (bottom panels) data. Excitatory neuron (*Slc17a7*), Inhibitory neuron (*Gad1*), Oligodendrocyte (*Plp1*), Astrocyte (*Aqp4*), and Endothelial cell (*Cldn5*). **B**, The spatial expression of cortex layer markers in the raw STARmap (top panels) and inferred by iSpatial (middle panels) compared with the ISH data from the ABA (bottom panels). Layer information: “L1 to L6”, the six cortical layers; “cc”, corpus callosum; “HPC”, hippocampus. **C**, The UMAP and spatial expression of *Tac2* and *Serpinb2* in the raw MERFISH (top panels) and inferred by iSpatial (middle panels) compared with the ISH data from the ABA (bottom panels).

In addition to the STARmap cortical dataset, a similar effect of iSpatial in correcting false-positive and false-negative expression on the MERFISH striatum data is also observed. Specifically, iSpatial removed most false-positive noise of known cell type-specific markers (**fig. S4A**). It also accurately predicted the expression and spatial pattern of genes not well detected by MERFISH, such as *Tac2*, *Serpinb2* and *Kctd4* (**Fig. 4C and fig. S4B**). Compared to the original MERFISH data, the spatial patterns inferred by iSpatial showed higher consistency with those determined by ISH (**Fig. 4C and fig. S4B**). For example, MERFISH indicates a broad distribution of *Serpinb2* across the striatum, but iSpatial suggested it was selectively expressed in a small group of cells located in the medial shell of NAc (**Fig. 4C**). ISH from ABA confirmed the accuracy of iSpatial’s prediction (**Fig. 4C**), suggesting that iSpatial is capable of reducing noise. Collectively, these results showed that iSpatial can reduce false-positive and false-negative signals in the original ST data from different tissues generated by different techniques.

### iSpatial enables whole-transcriptome level spatial analysis

One major goal of ST analysis is identifying SVGs, which are the molecular basis of structural/functional heterogeneity in different tissues. Since iSpatial could reliably infer genome-wide gene expression and their spatial locations, we sought to test whether iSpatial could augment the capability of certain ST dataset in detecting SVGs and spatial gene expression patterns. To this end, we applied iSpatial to the STARmap cortex dataset to identify SVGs. We found that iSpatial inferred data dramatically increased the number of detected SVGs (from 21 to 2,122, **fig. S5A**). Clustering analysis of the SVGs revealed six major spatial patterns (**fig. S5B**), which resemble the known layer organization of mouse cortex. Notably, even when we restricted the analysis to the target genes of STARmap, iSpatial still identified more SVGs (162 in inferred data and 21 in original data), likely due to the correction of false-positive and false-negative signals in the original data (see above). Indeed, most genes (73.2%) exhibited smaller p-values in the iSpatial inferred data compared to the original data when the same spatially variable test was applied (**fig. S5C**), suggesting higher SVGs detection power in the inferred data.

In addition to STARmap, we performed parallel analysis to evaluate iSpatial’s effect on SVG identification using the MERFISH striatum dataset (**Fig. 5A**) and observed a similar increase in the SVGs number and statistic power of spatially variable test. Specifically, the SVG number increased by > 20 folds (from 94 in the original data to 1,968 in the inferred data) (**fig. S5D**), and 71.9% of the target genes showed decreased p-values in spatially variable analysis (**fig. S5E**). Compared to cerebral cortex, the anatomic organization of striatum is more ambiguous and less well understood, although recent studies have suggested that distinct transcriptional features and cell types underlying its anatomic heterogeneity (*27, 28*). By unbiased clustering analysis of the SVGs obtained from the iSpatial inferred data, we identified 12 distinct spatial patterns of SVGs (**Fig. 5B**). Importantly, many of these patterns closely resemble the known anatomic subregions in the striatum. For example, the C12 cluster is mainly expressed in the dorsal striatum, while C1 and C4 clusters are highly enriched in the NAc (**Fig. 5B and 5C**). Additionally, C7 and C11 clusters correspond to the core region of the NAc, while the C2, C9 and C10 clusters represent the shell region (*29*) (**Fig. 5B and 5C**). Interestingly, the C8 cluster is specifically expressed in the medial shell (**Fig. 5B, 5C and fig. S5F**), a NAc subregion known to possess distinct anatomic and functional features (*30–32*). These results indicate that iSpatial can facilitate identification of biologically relevant spatial gene expression patterns.

**Fig. 5:**
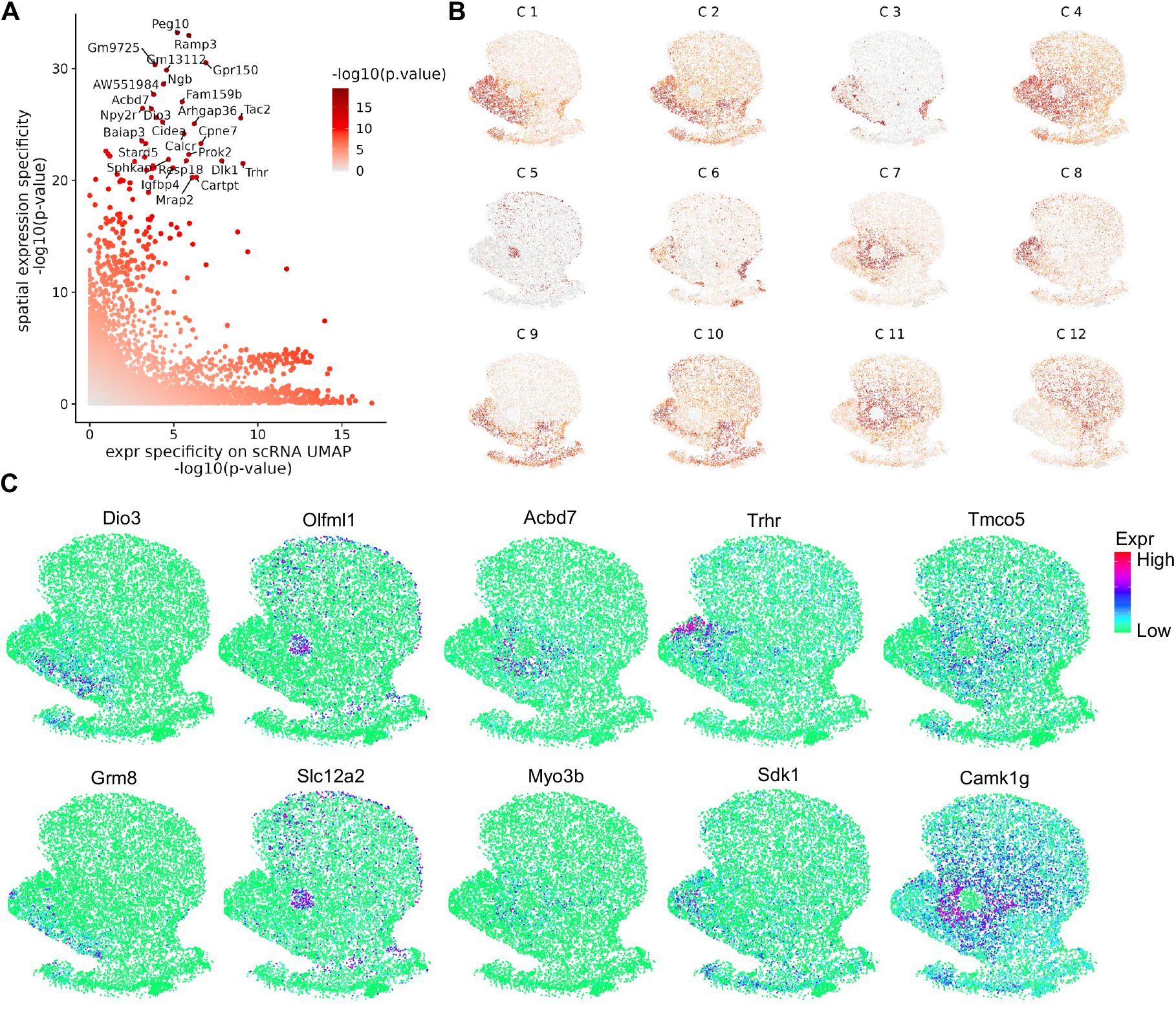
iSpatial enables whole-transcriptome level spatial analysis. **A**, Scatterplot showing genes with expression specificity in spatial location and UMAP projection of the corresponding scRNA-seq data. **B**, The 12 clusters of the spatial variable genes in mouse striatum. The plot is colored coded by the average gene expression in each cluster. **C**, Inferred spatial expression signals of representative genes in different clusters. C1: *Dio3* and *Grm8*, C5: *Olfml1* and *Slc12a2*, C7: *Acbd7* and *Myo3b*, C8: *Sdk1* and *Trhr*, C11: *Camk1g* and *Tmco5*.

### iSpatial improves analysis of spatial transcriptomic data from liver

Having demonstrated the utility of iSpatial in the analysis of ST data from different brain regions, we next sought to test iSpatial’s performance with ST data from other tissues. To this end, we analyzed a Vizgen MERFISH Mouse Liver Map dataset with 347 target genes included in the original data (https://vizgen.com/data-release-program/). By integrating the MERFISH data with a liver scRNA-seq dataset (*33*), iSpatial successfully inferred the expression of over 6,000 genes on average in each single cell (**Fig. 6A and 6B**), which increased by > 20 folds from the original data. Importantly, the inferred spatial patterns were largely consistent with established knowledge. For example, iSpatial predicted selective expression of *Slc1a2* and *Aldh1b1* in cells around the central vein (CV) and portal vein (PV), respectively (**Fig. 6C**). Similarly, *Cyp2e1* and *Cyp2f2* were predicted to be biased to CV and PV, but they have broader distribution than *Slc1a2* and *Aldh1b1* (**Fig. 6C**). All these spatial patterns were confirmed in previous studies (*34, 35*). We further generated UMAP based on iSpatial inferred expression profile and found a close correlation between cells’ positions on UMAP and their *in-situ* distribution along the CV-PV axis (**Fig. 6C and 6D**), revealing a gradient expression profile along the CV-PV axis. Notably, although Liger and Seurat can also reveal a similar gradient expression patten, a comparison among the three methods indicated that iSpatial achieved a higher specificity and accuracy, especially on genes with more spatially restricted expression patterns. For example, *Slc1a2* is selectively expressed in a monolayer surrounding the CV, which was accurately predicted by iSpatial (**Fig. 6C**), while Liger and Seurat revealed a more broader expression pattern (**fig. S6A and S6B**). Based on the observed relationship between gene expression and spatial location of liver cells, we calculated a CV score for each cell (see Methods) to reflect its relative position to CV/PV, with a high/low CV score indicating close to CV/PV, respectively. As expected, the CV score showed gradual increase from PV to CV in both the liver tissue and the UMAP space (**Fig. 6E and 6F**). Then all the genes in the inferred dataset (from iSpatial) were ranked according to their correlation with the CV score, which enabled us to systematically identify genes strongly related to cell’s spatial distribution (**Fig. 6G**). From this analysis, 141 and 692 genes with correlation to CV score > 0.3 or < −0.3 were predicted to be strongly enriched in cells close to CV or PV. In contrast, only 3 and 17 of these spatially variable genes were included in the original MERFISH dataset. As expected, many genes known to be biased to CV or PV were found, including *Gulo* and *Cyp2a5* enriched in cells adjacent to CV, *Cdh1* and *Etnppl* mainly expressed in cells around PV (**Fig. 6H**). Furthermore, by applying KEGG pathway analysis to CV and PV enriched genes, we found that they are involved in different biological functions (**Fig. 6I**). For example, the genes highly expressed around CV were enriched in “drug metabolism” and “PPAR signaling pathway”, while the genes highly expressed in PV were enriched in “protein processing in ER”, and “complement and coagulation cascades” (**Fig. 6I and fig. S6C**). These findings were consistent with previous reports (*34, 35*). Taken together, the above analyses demonstrate that iSpatial can overcome the limited target gene numbers of various ST analyses to the whole-transcriptome level with high accuracy in different tissues.

**Fig. 6:**
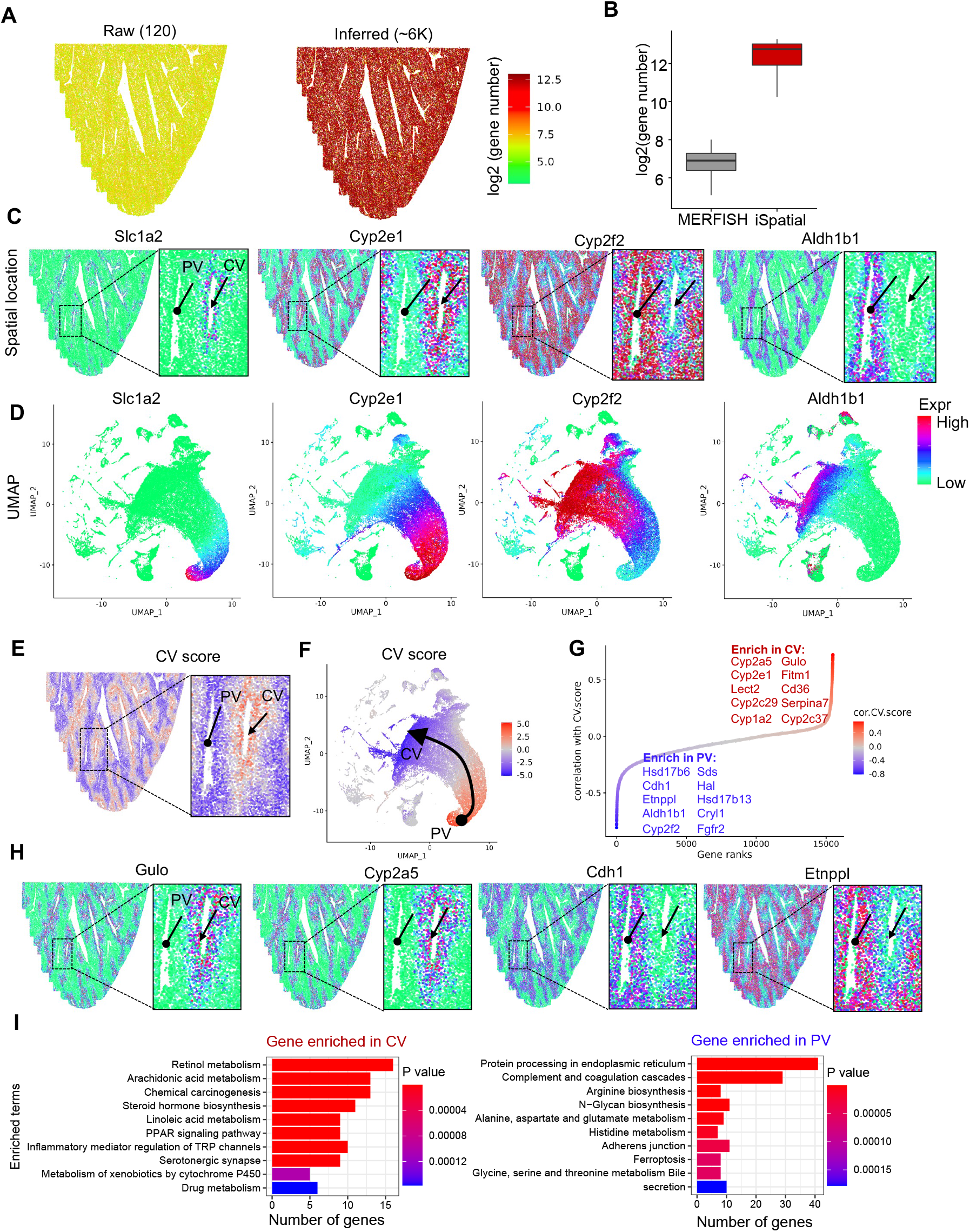
iSpatial infers the spatial expression patterns in liver. **A**, The numbers of detectable genes in each cell in raw merFISH (left) and after inferring by iSpatial (right). **B**, Boxplot showing the detectable gene numbers without or with interfering by iSpatial. **C**, Representative examples of genes exhibiting spatial patterns enriched in CV (*Slc1a2*, *Cyp2e1*) or PV (*Cyp2f2*, *Aldh1b1*). **D**, UMAPs showing the expression levels of the genes showed in (C). The UMAPs were generated with iSpatial inferred expression. **E**, The spatial location of each cell colored by CV score. **F**, The UMAP of all cells colored by CV score. **G**, The scatterplot showing the ranked correlations between gene expression level and CV score. The names of top 10 enriched genes in PV or CV are listed. **H**, Examples of genes selectively expressed nearby CV or PV. **I**, The top 10 enriched KEGG pathways of genes selectively expressed near CV or PV.

## DISCUSSION

Spatial transcriptomic assays simultaneously profile gene expression and their spatial location in tissue context, with the potential to unveil transcriptional features associated with tissue organization, cell-cell interaction, and region-specific physiological/pathological changes (*36*). Although sequencing and imaging-based ST techniques have been rapidly evolving (*5*), obtaining a genome-wide expression profile with single-cell spatial resolution is still challenging. To overcome this limitation, we developed a computational tool iSpatial to infer the genome-wide spatially resolved transcriptional information. iSpatial is especially useful for imaging-based ST analysis (such as MERFISH, seqFISH, STARmap), which in general has high detection efficiency and single-cell/subcellular spatial resolution, but is usually limited by the pre-defined gene targets. By integrating such kind of ST data with corresponding scRNA-seq profiles, the expression levels of untargeted genes could be inferred from scRNA-seq data, while the spatial information is directly inherited from the ST data, enabling high-resolution spatial analysis at the whole-transcriptome scale.

To ensure accurate expression imputation, it is critical to account for intrinsic noise in different original datasets. Specifically, high-throughput scRNA-seq has low capture efficiency on mRNA molecules, leading to a large proportion of zero counts for expressed genes (dropout). On the other hand, due to the variable performance of different gene probes, imaging-based ST analysis may generate both false-positive and false-negative signals. iSpatial includes an expression stabilization step, which borrows the information from cells with similar global expression pattern to minimize random noise in the original data (**fig. S7A**). In addition, iSpatial uses two-rounds integration to remove technology bias and batch effect on PCA space, allowing accurate integration of ST and scRNA-seq datasets. A comparative analysis indicated that two-round integration resulted in more accurate prediction than one-round integration (**fig. S7B**). As a result, iSpatial exhibits a significantly higher accuracy in both benchmark analysis with Slide-seq data and cross-validation with image-based ST data when compared with other existing imputation methods. Notably, iSpatial is not only able to faithfully predict the spatial expression of genes out of the original ST data, but also shows robust performance on reducing the false-positive/negative signals in the raw data. In both cortical STARmap and striatal MERFISH datasets, iSpatial is able to remove random expression of cell type-specific markers in non-relevant cell types and correct some non-specific background in the raw data (probably caused by poor performance of certain probes) to generate spatial expression patterns that are consistent with established knowledge. Since it is difficult to directly validate the detected mRNA sequences that generate the signals in imaging-based ST techniques, iSpatial provides a useful method to evaluate the performance of different probes.

We have tested iSpatial with multiple ST datasets generated from different tissues and techniques. In all the applications, iSpatial is able to expand spatial information from a pre-defined gene panel in the original ST data to the whole transcriptome, which renders several benefits for downstream analysis: First, it enables systematic identification of SVGs. In both the brain and liver datasets, we found that the number of SVGs increased from a few hundreds to several thousand after iSpatial imputation. Second, iSpatial enables the discovery of distinct spatial expression patterns across the tissue, which is achieved by spatial clustering analysis of SVGs. Importantly, many such unbiasedly identified expression patterns are biologically relevant. For example, we found that the SVG groups are organized into layer structure in the cortex and they exhibit core/shell enrichment in the striatum, indicating a tight relationship between gene expression and tissue organization. Third, iSpatial enables bioinformatic analysis requiring sufficient gene number or high statistical power, such as KEGG analysis, to be performed on SVGs or SVG subgroups. For example, by inferring the transcriptome-wide spatial pattern in liver, we found genes enriched in CV and PV are involved in distinct KEGG terms, suggesting a likely link between region-specific gene expression and function.

A potential limitation of iSpatial is that it requires corresponding ST and scRNA-seq data, which may not be always available. But given the rapid development of ST and scRNA-seq techniques, and ongoing effort in large single-cell consortium and spatial transcriptomics (*4, 37, 38*), we anticipate that iSpatial will be widely used to help understanding the molecular basis of structural and functional heterogeneity in complex tissues of diverse organs in normal and disease states.

To facilitate iSpatial implementation, we have made the iSpatial R package available at https://github.com/YiZhang-lab/iSpatial, where a tutorial on how to use iSpatial to integrate striatal scRNA-seq and MERFISH data to infer genome-wide spatial expression patterns can be found.

## MATERIALS AND METHODS

### iSpatial workflow

The workflow of iSpatial R-package contains the following steps: 1) Expression stabilization, which removes possible false positive and false negative gene expression signals at single cell level; 2) Integration. Two rounds of integration are performed to achieve accurate mapping of single cell RNA-seq data and spatial transcriptome data; 3) Infer expression according to the weighted k-nearest neighbors (KNN); 4) Downstream analysis. Detection of spatial variable genes and patterns.

### Expression stabilization

In the probe-based spatial transcriptome, we observed some cell type markers are not fully detected in corresponding cell cluster. However, these markers are indeed expressed across the cluster based on the scRNA-seq data. This indicates some probes have low binding affinities, which generates false negative (FN) signals. We also found some markers could be detected in some cell types which should not be expressed (random distributed in whole slice). This can be caused by nonspecific binding of some probes, which cause false positive (FP) signals. iSpatial try to remove these FN/FP signals to correct the expression in each cell. We assume that these FN/FP signals are randomly distributed. Thus, it can be corrected using the cells show similar global expression patterns, but don’t exhibit FN/FP on the same gene. To remove these FN and FP signals in each single cell *i*, we first find the K-nearest neighbors (*KNN_i,k_*) which most correlate with the cell *i* based on the global expression pattern.

*E_i_*: the vector of gene expression in cell *i*

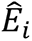: the vector of corrected gene expression in cell *i*

*KNN_i,k_*: {*KNN*_*i*,1_, *KNN*_*i*,2_, …, *KNN*_*i,k*_} : the set of k-nearest neighbor of cell *i*

Then, we correct the gene expression 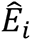 by the expression of KNNs:

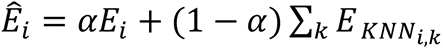

where *α* controls the weight of expression from cell *i* and neighbors. By default, we set *α* = 0.5.

### Integration of spatial transcriptomic data and scRNA-seq data

To achieve accurate integration of spatial transcriptomic data and scRNA-seq data, iSpatial uses a two-round integration approach. In the first round, we adopted a reciprocal principal component analysis (RPCA) method from Seurat. At this step, the PCA spaces are calculated in both datasets. Then, one dataset is projected onto the other’s PCA space and constrain the cell anchors with the same mutual neighbors. We use the “FindIntegrationAnchors” function with the parameter reduction = “rpca” in Seurat (version 4.0.5) to find the anchors and use “IntegrateData” to get the integrated data. For the second-round integration, we project the cells from both spatial transcriptome and scRNA-seq into a shared PCA embedding. Then an iterative clustering method is used to remove the technology bias and batch effect on PCA space, which get harmonious PCA embeddings for both datasets. Harmony (version 0.1.0) is then used to generate normalized PCA embeddings in this step.

### Infer spatial expression

After integrating two datasets into one reduced dimension space, iSpatial uses a weighted KNN approach to infer the expression of nontargeted genes. For each cell *t* in spatial transcriptome data, iSpatial searches the K-nearest neighbors (*KNN_t,k_*).

*t*: cell in spatial transcriptome data

*c*: cell in scRNA-seq data

*KNN_t,k_*: {*KNN*_*t*,1_,*KNN*_*t*,2_, …, *KNN*_*t,k*_} : the set of k-nearest neighbor of cell *t*

Then, *KNN_t,k_* are restricted to the cells from scRNA-seq data. Because the cells from scRNA-seq data contain the expression information of the whole transcriptome.

*KNN*′*_t,k_*: *KNN_t,k_* ∈ *c*

The final inferred expression 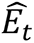 of cell *t* is calculated by the expression of cell *t* itself and the expression of *KNN*′*_t,k_*:

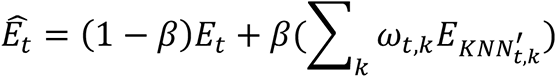

where *ω* is the weight of each neighbor *k* of cell *t*. For the genes targeted in cell *t*, *β* balance the expression from spatial transcriptome and scRNA-seq. *β* ∈ [0,1]. The weights *ω* between cell *t* and its neighbor *KNN*′_*t*_ are defined by the normalized transcriptional distance *d_t,k_*. Here, iSpatial use 1 - Pearson’s correlation coefficient to measure the distance:

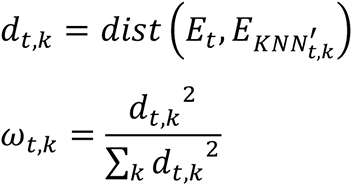

And,

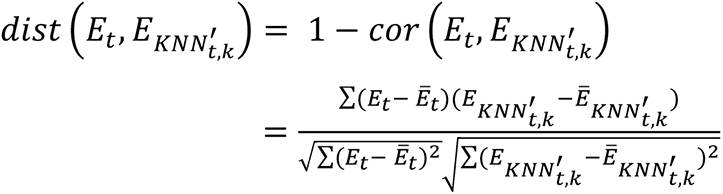

### Identifying spatially variable genes

To identify significant spatially variable genes, the x and y axes of the spatial location are evenly divided into *n* bins, then the spatial location is further grided into *n* × *n* grids. For each gene *j*, we calculate the average expression values *E_j_* over the *n*^2^ grids. We then randomly sample the spatial location of each cell and calculate the average expression *E*′*_j_* over the randomly sampled *n*^2^ grids. If a gene *j* has no specific spatial expression pattern, the distribution of observed *E_j_* should not be different from that of random *E*′*_j_*. On the contrary, if a gene has strong spatial expression pattern, the distribution *E_j_* should be significantly different from *E*′*_j_*. Thus, whether a gene exhibits spatial expression pattern depends on whether there is a difference between the distribution of *E_j_* and *E*′*_j_*. Here, we apply a non-parametric two-sides Mann–Whitney U test to determine the difference between *E_j_* and *E*′*_j_*. We also offer the Kolmogorov-Smirnov test to test the distribution differences.

Some studies found that a gene with spatial expression pattern always display a specific expression bias on scRNA-seq UMAP/tSNE projection. iSpatial also integrates scRNA-seq information into spatial variable gene detection. If a gene not only displays spatial expression pattern on spatial transcriptome data but also exhibits expression specificity on scRNA-seq UMAP/tSNE projection, this gene has a higher confidence of spatial expression pattern. Similar to the detection of spatial expression gene on spatial location, iSpatial uses the same method to detect whether a gene displays a specific expression location on scRNA-seq UMAP/tSNE. To integrate these two lines of information, the final p value is equal to the p value from the spatial transcriptomic data multiplied by the p value from the scRNA-seq data. Then the adjusted p values are calculated to control the false discovery for multiple comparisons.

### Spatially variable gene clustering

The genes with spatial expression patterns could be grouped into clusters. For each gene, iSpatial captures the spatial expression features according to the average expression value over the *n*^%^grids, which was described before. Based on these features, Pearson’s correlation coefficients are calculated for pairs of genes. Then, distances among genes are measured by 1 minus Pearson’s correlation coefficients. Finally, Hierarchical clustering is performed using “hclust” in R to spatial variable genes. “cutree” function from R is used to group these genes into desired number of groups.

### Single-cell RNA-seq data processing

The initial gene×cell matrix for each study was downloaded according to the original papers. The expression matrix was then transferred into Seurat object for downstream analysis. The raw counts of gene expression profile of each cell were normalized to 10,000 counts and natural log transformed using the Seurat function “NormalizeData”. To generate UMAP, we used standard pipeline from Seuart. In short, “FindVariableFeatures” was used to identify top variable genes, “ScaleData” was used to scale and center these genes in the data, then PCA was performed by “RunPCA” based on the selected features. Finally, the top 30 principal components from PCA were used to generate the UMAP projection by “RunUMAP” with the parameters “dims=1:30”.

### Mouse hippocampus data processing

The Slide-seq V2 of mouse hippocampus was download from the Broad Institute single cell portal website (https://singlecell.broadinstitute.org/single_cell/study/SCP815). Here, we only use the “Puck_200115_08” dataset (*24*). This dataset contains two files, the raw expression matrix and the barcode locations. The data processing of Slide-seq V2 followed the same procedure as that of single cell RNA-seq. The main difference is that Slide-seq contains spatial location of each cell. According to the vignettes of Seurat, the coordinate of each cell is stored as a “SlideSeq” class in Seurat object. For the scRNA-seq data of mouse hippocampus, we used a published dataset (*1*). To facilitate analysis, we used a pre-processed Seurat object (https://www.dropbox.com/s/cs6pii5my4p3ke3/mouse_hippocampus_reference.rds?dl=0) offered by the Satija Lab.

### Mouse cortex data processing

The STARmap data of mouse visual cortex was download from the STARmap resource website (https://www.starmapresources.org/data). We chose the dataset “20180505_BY3_1kgenes” which profiles 1020 genes. The expression matrix data “cell_barcode_count.csv” was imported to Seurat object. The spatial location coordinate of each cell was extracted from “labels.npz” according to the method provide by the original paper (https://github.com/weallen/STARmap). The cell coordinates were integrated into Seurat object. The single cell RNA-seq was download from the Allen Brain Atlas (https://portal.brain-map.org/atlases-and-data/rnaseq/mouse-v1-and-alm-smart-seq). The “mouse_VISp_2018-06-14_exon-matrix.csv” file was used to generate the expression profile. Low-quality cells were removed according to the meta data “mouse_VISp_2018-06-14_samples-columns.csv”.

### Mouse striatum data processing

The mouse striatum merFISH data was processed and normalized as described in the original paper (*27*). Briefly, the expression of each cell was normalized by cell size and total RNA counts. Then, the log-transformed data was applied to the expression matrix. In this manuscript, we used the data from a representative slice (slice 10). We used the pre-process Seurat object data in https://www.dropbox.com/s/ghkcovukgtctm76/NA_merFISH.RDS?dl=0. A down-sample version of this data are provided by iSpatial package as a test dataset. After installing the R package, this command “data(NA_merFISH)” could load the merFISH data into R environment. The scRNA-seq data of mouse striatum was downloaded from GEO under accession GSE118020. To speeding up the analysis, we down-sampled the full dataset to 10,000 cells using R function “sample”. Then this data was used to infer the spatial expression of all genes.

### Mouse liver data processing

The mouse liver merFISH data is download from Vizgen MERFISH Mouse Liver Map (https://vizgen.com/data-release-program/), which targets 347 genes. This dataset contains multiply slices. We only used slice 3 from the first replicate in this manuscript. The merFISH data processing is the same as described above. The matched scRNA-seq data was download from GEO under the accession GSE166504 (*33*). The original data set not only contains wild-type healthy livers but also livers with nonalcoholic fatty liver disease (NAFLD). Three healthy samples (Hepatocyte_Chow_Animal1_Capture1, Hepatocyte_Chow_Animal2_Capture1, Hepatocyte_Chow_Animal3_Capture1) were used in this analysis.

### CV score calculation

To measure the distance between each cell and central vein (CV), we calculated a CV score of each cell based on well-known CV markers (*Slc1a2*, *Cyp2e1*) and PV markers (*Cyp2f2*, *Aldh1b1*). For each cell, the CV score was defined by the mean expression level of *Slc1a2*, *Cyp2e1* minus mean expression level of *Cyp2f2*, *Aldh1b1*. A positive value means more likely that the cell is located near the CV. On the contrary, a negative value indicates that the cell is located near the PV. After defining the CV score, we could then identify the genes specifically expressed in CV/PV. For each gene, we calculated the Pearson’s correlation between the corresponding expression value and CV score across all cells. A positive correlation coefficient represents gene preferentially expressed in CV, and a negative one represents gene preferentially expressed in PV. According to the distribution of the correlation coefficient among all genes, we manually chose > 0.3 or < −0.3 as cutoffs to define genes with most CV/PV bias.

### Visualize the spatial expression

Visualization of gene spatial expression was achieved using the “SpatialFeaturePlot” in the Seurat package. iSpatial provides a function “spatial_signature_plot” for spatial visualization of the mean expression of a group of genes.

### Benchmark methods

We chose two popular methods for comparison: Seurat (version 4.0.5) and rliger (version 1.0.0). The Seurat provides a “canonical correlation analysis” (CCA) method to integrate the spatial transcriptome and scRNA-seq. Here we used the function “FindTransferAnchors” with parameter “reduction = “cca”” to identify the integrated anchors. And then genes expression values from scRNA-seq data were transferred to spatial transcriptome data using “TransferData” with the parameter “weight.reduction = “cca””. Different from Seurat, Liger uses an integrative non-negative matrix factorization (iNMF) method to integrate the datasets, which is embedded in the function “optimizeALS”. For imputing the non-targeted genes by rliger, we performed following steps according to the rliger vignettes: “createLiger”, “normalize”, “scaleNotCenter”, “optimizeALS”, and “quantile_norm”. Finally, “imputeKNN” was used to search the nearest neighbors and impute the non-targeted genes.

### Performance evaluation

The Slide-seq V2 data contains a total of 23,264 genes. We randomly sampled 3,000 genes as the training dataset and used others as validation. After performing different methods to infer the spatial expression of all genes, we conducted the comparisons between the inferred expression values and the raw expression values. At cell level, we calculated the Pearson’s correlation coefficient of each gene across all cells, then compared the overall differences of correlation coefficients among all genes of different methods. At spatial expression level, we grided the spatial location into 50×50 bricks. Then, we calculated the mean expression in each brick and compared the differences between raw expression and inferred one using 50×50 bricks as features. In addition, we compared the detection accuracies of spatial variable genes among three methods. The spatially variable genes were also called using validation dataset as the ground truth. Receiver operating characteristic (ROC) curves and area under the curve (AUC) were performed by R package pROC (version 1.18.0) (*39*).

### Ten-fold cross validation

For the datasets other than Slide-seq which we don’t have the ground truth, we used 10-fold cross validation to evaluate the prediction performance. For the targeted genes in spatial transcriptome, we randomly separated it into ten groups. Each time we chose nine groups to infer the expression and used the left one to validate. After ten rounds of inferring and validation, every gene was used to validate the prediction performance. Then, correlation of gene level and spatial expression level were calculated as described above.

## Data and Software Availability

iSpatial is available as an R package on github (https://github.com/YiZhang-lab/iSpatial).

## Acknowledgements

We thank Dr. Aritra Bhattacherjee for some initial discussion and for critical reading of the manuscript. This project was supported by 1R01DA042283, 1R01DA050589, and HHMI. R.C. is partially supported by the 2020 NARSAD Young Investigator Grant from the Brain and Behavior Research Foundation. Y.Z. is an investigator of the Howard Hughes Medical Institute.

## Author Contributions

Y.Z. and C. Z. conceived the project. C.Z. developed the iSpatial package and performed the data analysis. C.Z., R.C., and Y.Z. designed the analyses, interpreted the results, and wrote the manuscript.

## Competing Interests

The authors declare no competing interests.

## Supplementary Figure

**fig. S1:**
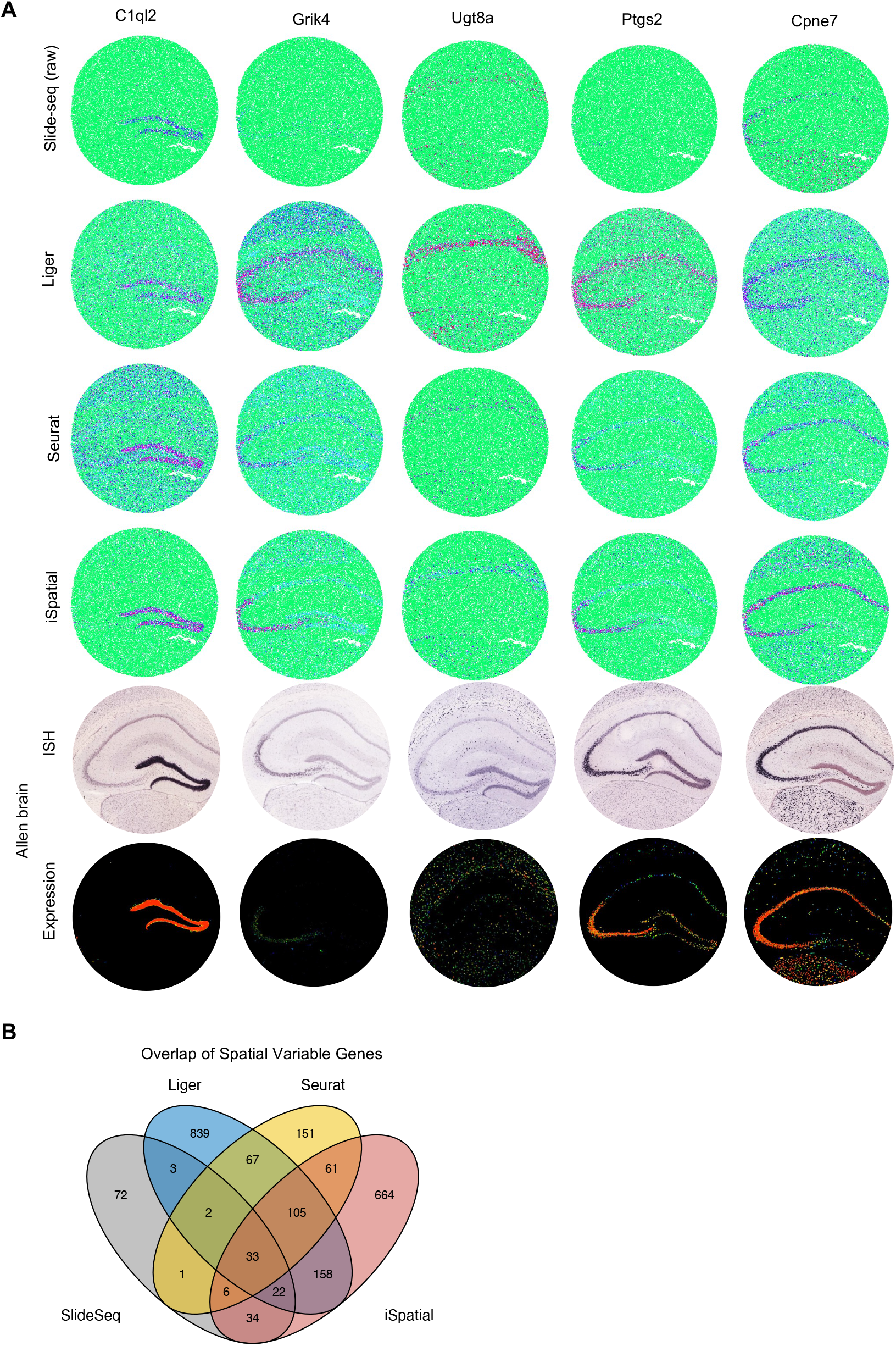
Benchmarking the performance of iSpatial in inferring genome-wide spatial transriptome. **A**, Representative examples showing the performance of Liger, Seurat and iSpatial in inferring spatial transcriptome. **B**, The overlaps of the spatial variable genes detected by different methods.

**fig. S2:**
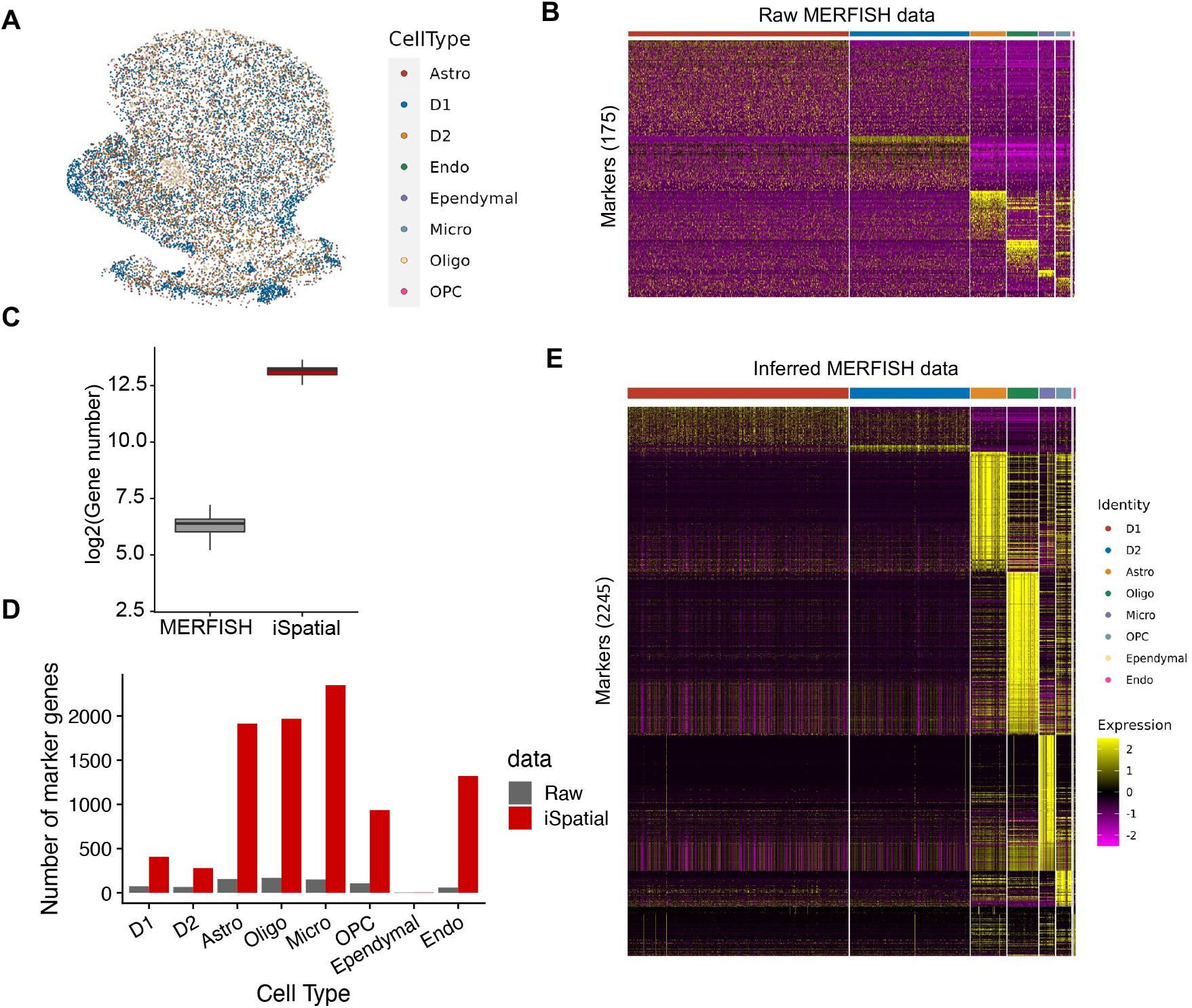
iSpatial is applicable to MERFISH dataset. **A**, Spatial distribution of the cell types detected by MERFISH. **B**, The heatmap of the 175 cell type markers detected in the raw MERFISH data. **C**, The numbers of detected genes in raw MERFISH data and after interfering by iSpatial. **D**, The number of marker genes in each cell type detected in raw and iSpatial inferred data. **E**, The heatmap of all maker genes detected after inferring by iSpatial.

**fig. S3:**
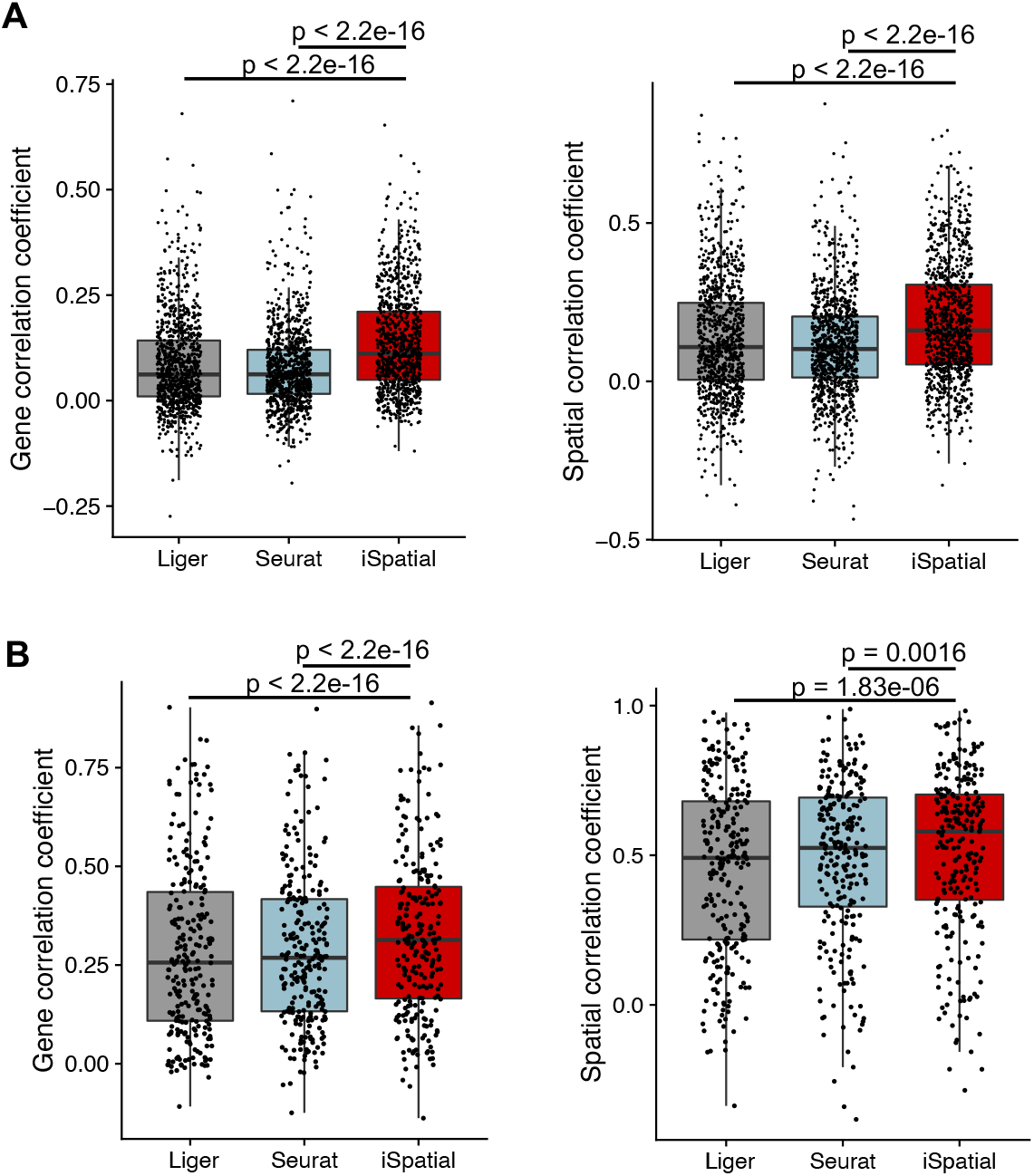
Ten-fold cross validation on the performance of the three methods in mouse cortex. **(A) and striatum (B)**. Left panels: gene expression correlation between raw expression data and inferred data. Right panels: spatial expression correlation between raw expression data and inferred data.

**fig. S4:**
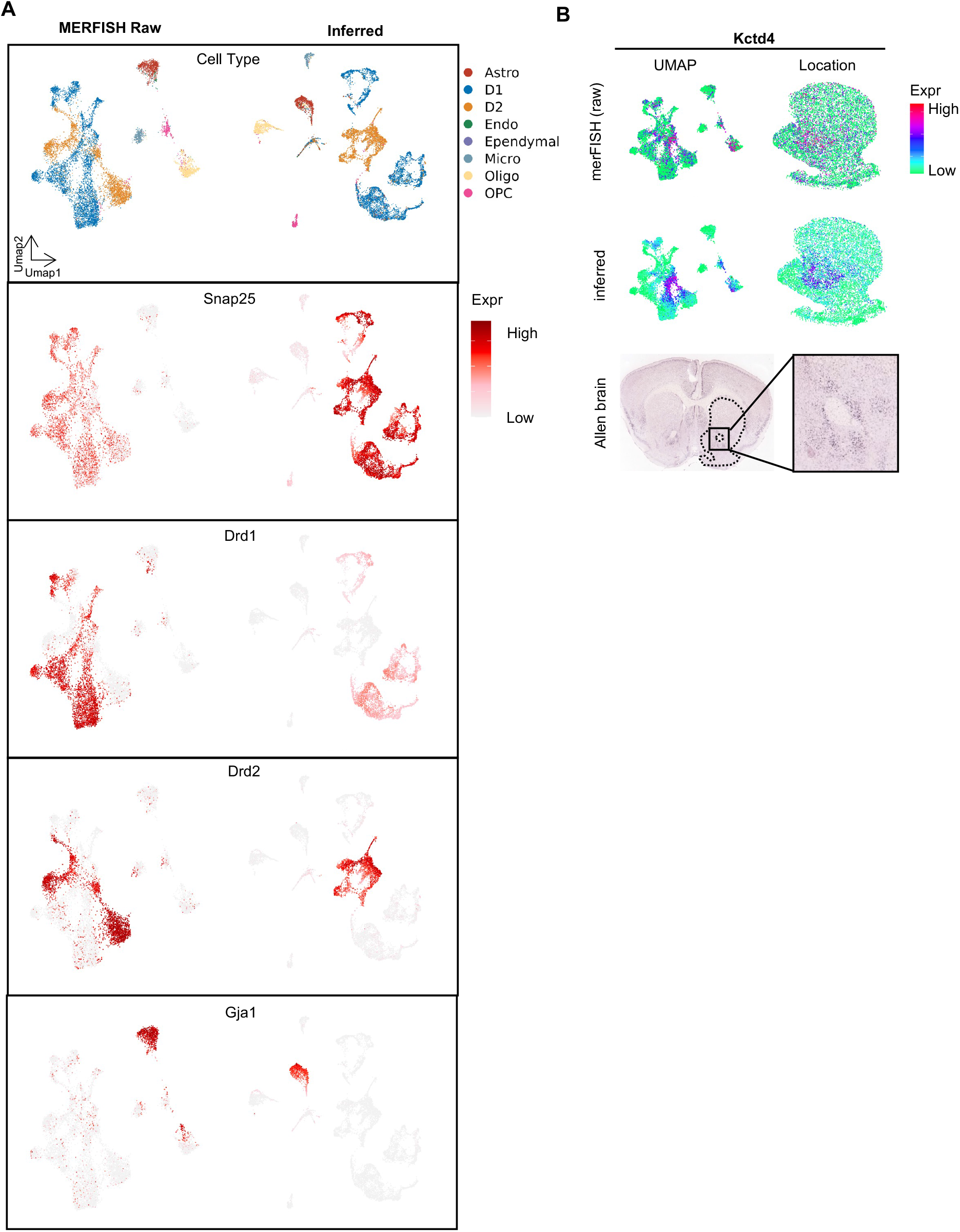
iSpatial can reduce false-positive and false-negative signals in the original MERFISH data. **A**, UMAPs showing the expression of representative cell type marker genes. The UMAPs are generated from raw or iSpatial inferred expression profiles, respectively. Marker genes for neuron (*Snap25*), D1 medium spiny neuron (*Drd1*), D2 medium spiny neuron (*Drd2*), and astrocytes (*Gja1*) are presented. **B**, The UMAP and spatial expression of *Kctd4* in the raw MERFISH (top panels) and inferred by iSpatial (middle panels) compared with the ISH data from the ABA (bottom panel).

**fig. S5:**
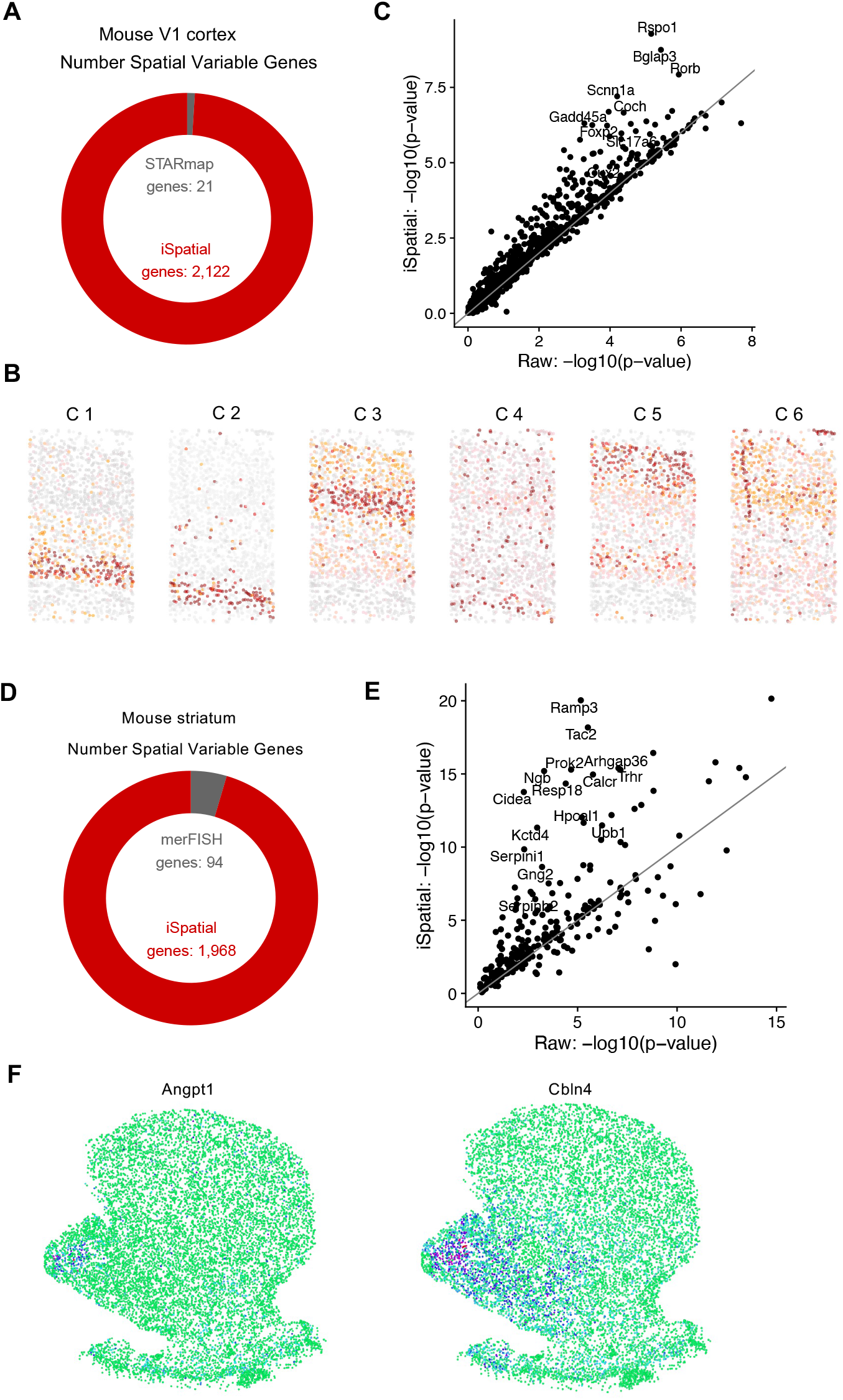
iSpatial detects spatial variable genes in mouse cortex (A-C) and striatum (D-F). **A**, The number of spatial variable genes in mouse V1 cortex detected by STARmap and after iSpatial interferring. **B**, The layer distribution of the 6 clusters spatial variable genes. The plot is colored by the average expression level of the genes in each cluster. **C**, Scatterplot showing the p-values of genes detected as spatial variable genes in STARmap raw data and iSpatial inferred data. **D**, The number of spatial variable genes in mouse striatum detected by MERFISH and after iSpatial interfering. **E**, Scatterplot showing the p-values of genes detected as spatial variable genes in MERFISH raw data and iSpatial inferred data. **F**, Inferred spatial expression signals of representative genes in the C8 cluster.

**fig. S6:**
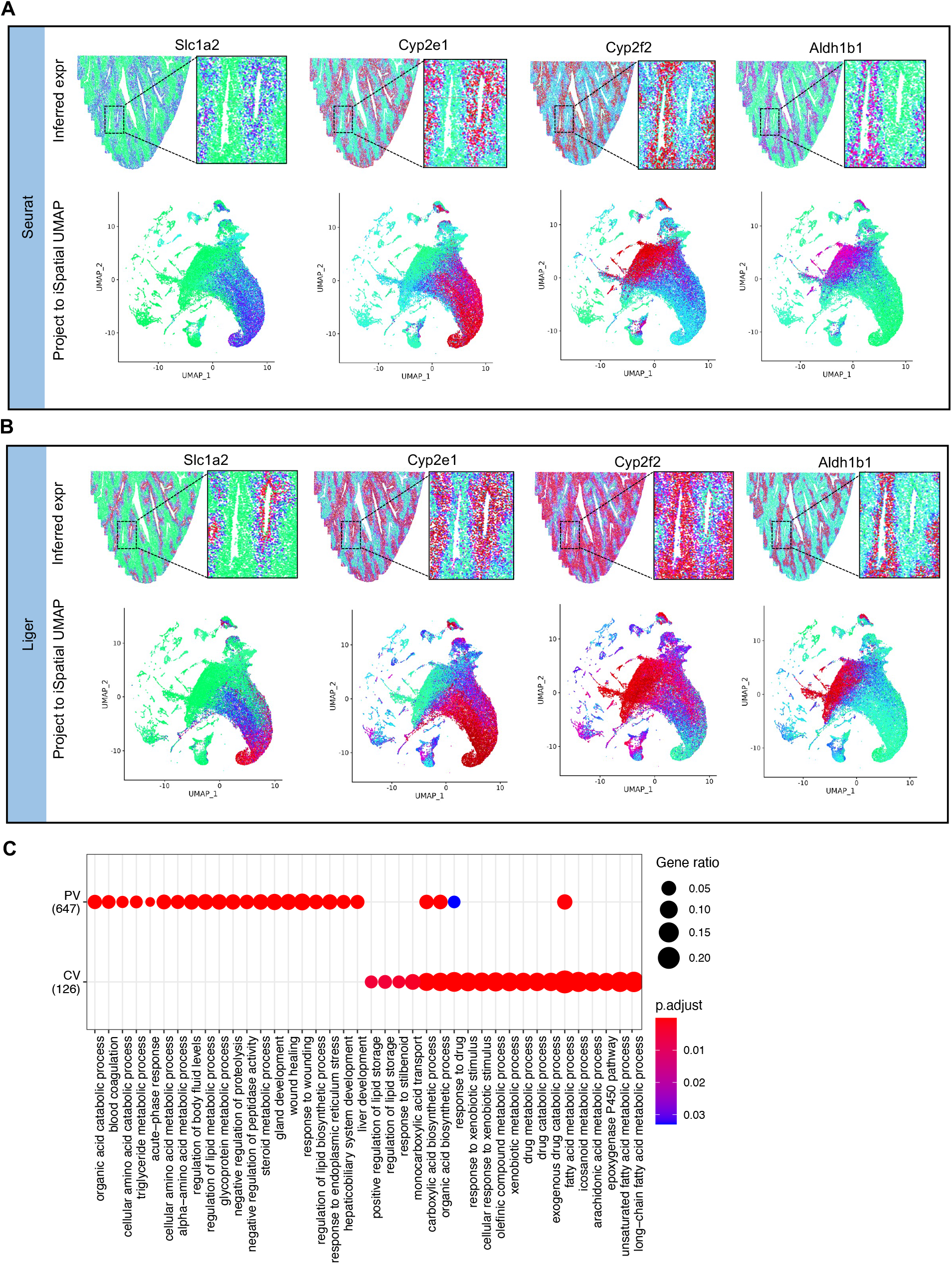
iSpatial infers the spatial expression patterns in liver. **A, B**, Examples of spatial expressed genes enriched in CV (*Slc1a2*, *Cyp2e1*) or PV (*Cyp2f2*, *Aldh1b1*), inferred by Seurat (A) and Liger (B). The upper panels show the spatial expression patterns. The bottom panels show the inferred genes’ expression by the indicated method projects on the iSpatial UMAP. **C**, The top 20 enriched Gene Ontology terms of genes selectively expressed near CV or PV.

**fig. S7:**
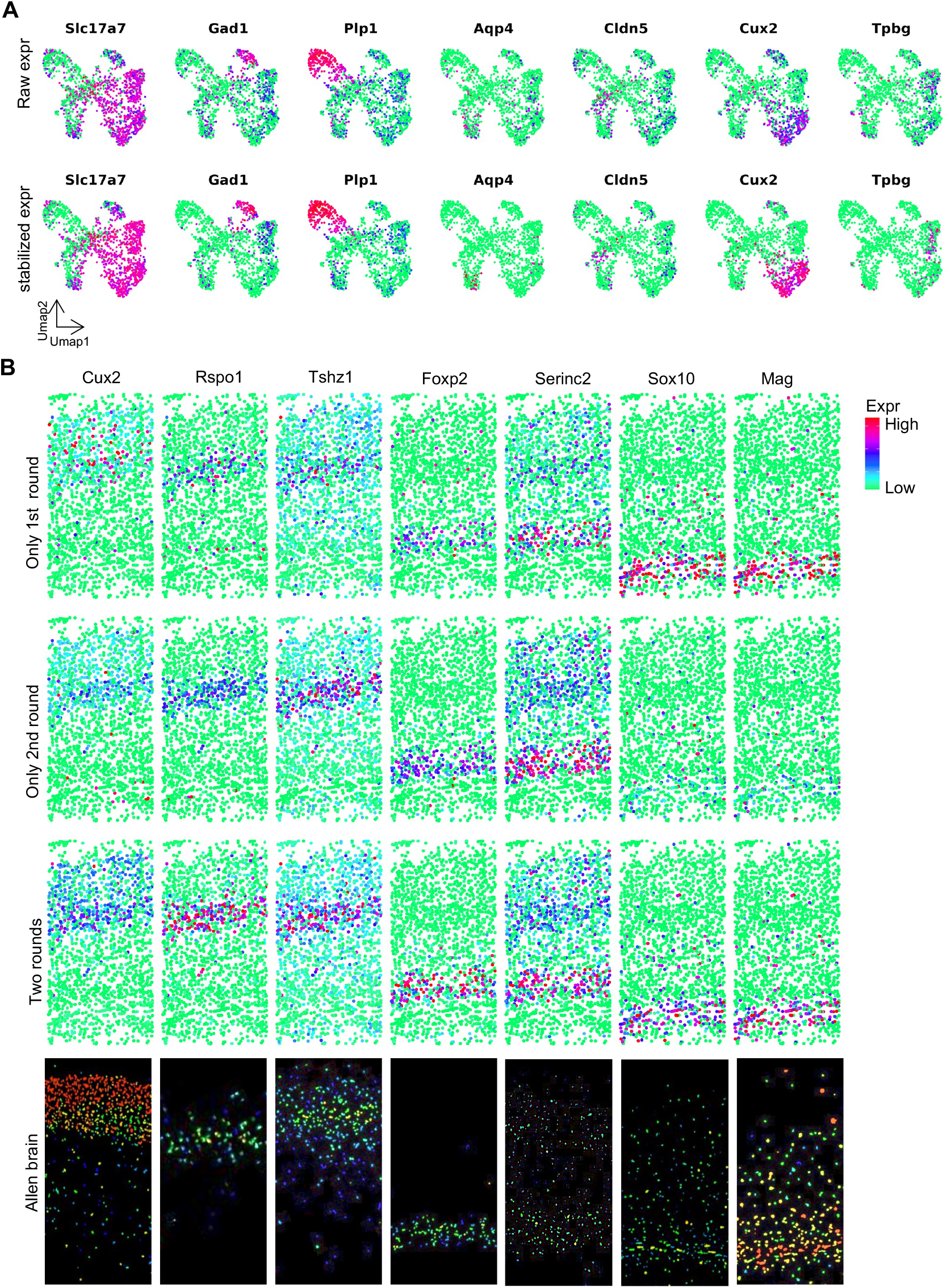
Stabilization and two-rounds integration steps contribute to the performance of iSpatial. **A**, UMAPs showing the expression signals of representative markers before (top panels) and after (bottom panels) expression stabilization in mouse cortex data. **B**, The spatial expression of the representative genes of the mouse cortex raw STARmap data inferred by one round (top two rows) or two rounds (third row) of integration by iSpatial compared with the ISH data from the ABA (bottom row). Reciprocal PCA is used for the first-round integration, and Harmony is used for the second-round integration.

## Notes

### Competing Interest Statement

The authors have declared no competing interest.

### Summary of Updates

Figure 1 revised

